# Context-Dependent Synergism and Antagonism between IGF2BP1/3 and METTL3/14 in m6A-mediated mRNA regulation

**DOI:** 10.1101/2025.05.04.652102

**Authors:** Ruchi Bhardwaj, Harsh Bhakhri, Jayanth Kumar Palanichamy

**Affiliations:** Department of Biochemistry, All India Institute of Medical Sciences, New Delhi, India

**Keywords:** RNA-Binding proteins, m6A machinery, readers, writers, mRNA stability

## Abstract

N-6 Methyl Adenosine (m6A) methylation is primarily found in the 3’-UTRs of mRNAs, near the stop codon, and at consensus sequence RRACH. The methylation reaction is catalyzed by RNA methyltransferases METTL3 and METTL14, known as writers. Readers recognize the modified mRNA, which influences mRNA stability and translation. Insulin-like growth factor-2 binding proteins 1 and 3 (IGF2BP1 and IGF2BP3) are known m6A readers, promoting mRNA stabilization. Erasers like ALKBH5 and FTO remove m6A modifications. Dysregulation of m6A machinery has been implicated in cancer progression.

We analyzed the expression patterns of writers, erasers, and readers (WERs) in multiple public datasets, including NCBI-GEO, TCGA, TARGET, and normal tissue expression data from GTEx. Our findings revealed widespread dysregulation of WERs across various cancers. To investigate whether IGF2BP1/3 and METTL3/14 function synergistically in mRNA stabilization, we identified direct mRNA targets using intersection analyses of IGF2BP1/3 eCLIP and METTL3/14 knockout (KO) datasets on Galaxy server. This analysis identified METTL14-dependent targets (*KDM3B, DYNLL1, CNOT1, RPL29*) and METTL3-dependent targets (*SREBF2, HNF4A, GNA11*), all bound by IGF2BP1/3.

To validate these interactions, we cloned 3’-UTRs of these targets downstream of luciferase reporter and assessed mRNA stability following IGF2BP1/3 and METTL3/14 overexpression. Luciferase activity increased for *RPL29, DYNLL1, SREBF2*, and *CNOT1* upon co-expression, indicating IGF2BP1/3-mediated mRNA stabilization in an m6A-dependent manner. In contrast, *KDM3B, HNF4A,* and *GNA11* exhibited reduced luciferase activity, suggesting destabilization.

Our study provides novel evidence that m6A readers and writers exhibit synergistic and antagonistic interactions in a target-dependent manner, underscoring the complexity of m6A-mediated gene regulation in oncogenesis.

## Introduction

RNA undergoes many chemical modifications, including methylation, acetylation, and phosphorylation. Changes in these modifications can lead to dysregulated gene expression. These modifications regulate various biological processes, including mRNA stability, localization, mRNA trafficking, and translation.

N-6-methyl adenosine (m6A) and 5-methyl cytosine (m5C) methylation are among the most common chemical modifications in eukaryotic mRNA (Q. Li et al. 2017). m6A methylation is catalyzed by the methyltransferases METTL3 and METTL14, commonly referred to as writers. The m6A mark is recognized by many RNA-binding proteins like YTHDF2, YTHDF1, IGF2BP3, and IGF2BP1, collectively known as readers. These readers, along with writers, regulate mRNA stability and translation, impacting gene expression.

m6A is a reversible modification due to the action of demethylases such as ALKBH5 and FTO, which can remove the methyl group from mRNA (Balacco and Soller 2019; Wu et al. 2016). m6A is the most prevalent modification in mammalian RNA that occurs at an estimated frequency of 3−5 sites per transcript. The sites enriched with m6A were mapped globally by pulldown and high-throughput sequencing approaches and were found to be distributed at specific sites in the mRNA. The m6A sites are enriched within a consensus sequence RRACH (R = G/A; H = A/C/U). Most m6A sites are found in the 5′ UTR, near the stop codon in the 3′ UTR, and within long exons (Wu et al. 2016; Vu et al. 2017). m6A is widely implicated in many cancers, like pancreatic, breast, liver, blood, and gastrointestinal cancers (Shen et al. 2023; Maity and Das 2016; Deng et al. 2018). Evidence suggests that METTL3 and METTL14 play a crucial role in acute myeloid leukemia (AML) development (Liu et al. 2014). METTL3 acts as an oncogenic factor in AML, gastric cancer, liver cancer, pancreatic cancer, colorectal cancer, and glioblastoma (Zeng et al. 2020). Additionally, METTL3-mediated m6A modification promotes esophageal cancer progression via the Notch signaling pathway (Han et al. 2021).

Insulin-like growth factor 2 binding proteins (IGF2BPs) are the readers that preferentially bind m6A at imprinted GGAC sites in the mRNAs and enhance their stability (H. Huang et al. 2018) and translation (Zhu, Hong, and Ling 2023). Structurally, IGF2BPs contain RNA recognition motifs (RRMs) and heterogeneous nuclear ribonucleoprotein (hnRNP) K-homology (KH) domains, which are critical for m6A binding and target recognition (Bell et al. 2013). IGF2BPs have been shown to stabilize key oncogenic transcripts like *MYC*, thereby promoting cancer progression (X. Huang et al. 2018). IGF2BP1 binds to m6A-modified targets in a sequence-specific manner, recognizing the GGm6AC consensus sequence (Bell et al. 2013). This binding stabilizes the mRNA of the target and further facilitates its expression. IGF2BP1 is overexpressed in many cancers, like breast cancer, sarcoma, gastrointestinal cancers, lymphoma, and leukemia (AML and B-ALL) (X. Huang et al. 2018; Gu et al. 2014). IGF2BP1 binds to the mRNA of *C-MYC, PTEN, IGF2, CD44,* and stabilizes their transcripts, promoting tumorigenesis. IGF2BP1 is highly expressed in *ETV6-RUNX1*-positive B-ALL patients and synergizes with ETV6:RUNX1, whereas IGF2BP3 is overexpressed in MLL-rearranged B-ALL (Sharma et al. 2021; 2023; Palanichamy et al. 2016).

IGF2BP3 contributes to glioblastoma progression by binding to the 5′-untranslated regions of IGF-2 mRNA and increasing its translation (Suvasini et al. 2011). IGF2BP3 binds to its downstream targets, *C-MYC* and *CDK6,* at the 3’UTRs of and stabilizes their mRNA, leading to an increased expression of these oncogenes. Functional studies indicate that IGF2BP3 is highly expressed in leukemic cell lines, and its knockdown leads to reduced cellular proliferation (Palanichamy et al. 2016).

The cooperative function of m6A writers and readers’ work has been demonstrated in numerous studies. The enzyme spermine synthase is a common target of METTL3 and IGF2BP3, which play an essential role in the progression of pancreatic cancer. The METTL3-IGF2BP3 complex stabilizes the mRNA of spermine synthase, which leads to a progression of pancreatic cancer by regulating phosphorylation in the PI3K-AKT pathway (Guo et al. 2022). The METTL3-IGF2BP3 axis has also been involved in inhibiting antitumor activity by stabilizing the mRNA of *PD-L1* in breast cancer (Wan et al. 2022).

Despite increasing evidence supporting the role of individual readers and writers, few studies have explored their combined effects. Our study aimed to investigate how IGF2BP1/3 and METTL3/14 cooperatively influence mRNA stability and translation of oncogenic targets. We examined the stability of transcripts targeted by the readers, IGF2BP1/3, and writers, METTL3/14, and validated their impact using a luciferase assay.

## Results

### Dysregulated expression profile of m6A WERs (writers, erasers, and readers) across various cancers

To investigate the gene expression profile of m6A machinery components—writers, erasers, and readers (WERs) across cancer types, we performed an integrative analysis using publicly available datasets. Gene expression data were retrieved from NCBI-GEO, TCGA, and TARGET pan-cancer studies, and normal tissue expression profiles were obtained from the GTEx repository. A comparative gene expression matrix was generated for key m6A machinery genes. Analysis revealed that most m6A-related genes were significantly dysregulated across various cancers compared to corresponding normal tissues (Figure 1A). Notably, METTL3 and METTL14 (writers), along with IGF2BP1 and IGF2BP3 (readers), showed elevated expression in acute myeloid leukemia (AML), diffuse large B-cell lymphoma (DLBC), cholangiocarcinoma (CHOL), kidney chromophobe (KICH), and testicular germ cell tumors (TCGT) in the TCGA and TARGET datasets (Figures 1A, S1). An increased expression of METTL3 and METTL14 (writers) and IGF2BP1/3 (readers) was observed in the AML, DLBC, CHOL, KICH, and TCGT TCGA/TARGET studies (Figure 1a, S1).

**Figure 1:**
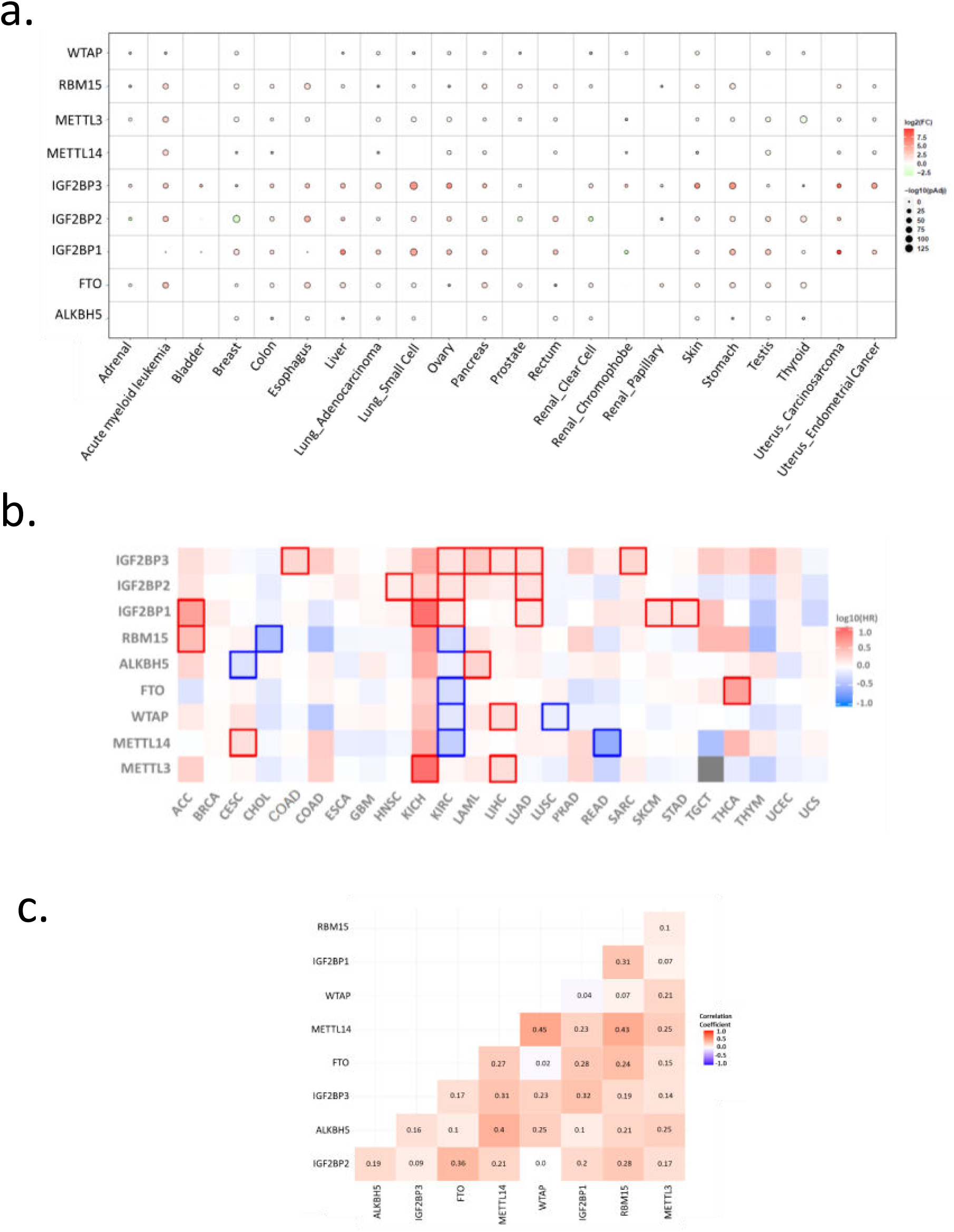
Gene expression analysis of the m6A writers, erasers, and readers (WERs) from the NCBI-GEO, TCGA, and TARGET pan-cancer studies compared to the GTEx repository. **a)** Pan-cancer gene expression dot-matrix plot for the WERs. The log2FC values represent tumor/normal RNA-Seq data, the red color represents higher tumor-tissue expression, and the green indicates higher normal-tissue expression. The size of the circle represents the negative log of the adjusted P values. **b)** Pan-cancer survival correlation with WERs expression. Log10 (HR) represents the hazard ratios for overall survival calculated for each gene between the high and low expression groups. Red indicates higher gene expression correlates with poorer survival (HR > 1), and blue indicates better survival (HR < 1). A solid box represents a Log10 (HR) value with a significant p-value. **c)** Gene expression correlation matrix of the WERs in tumor-tissues from the AML TARGET study. The correlation coefficient value represents the Spearman correlation coefficient, with the red color representing a positive correlation coefficient and the blue color representing a negative correlation coefficient.

To understand the clinical relevance of these alterations, we analyzed their association with patient survival across TCGA cohorts. The hazard ratio elevated expression of METTL3, METTL14, IGF2BP1, and IGF2BP3 correlated with poorer overall survival in AML, liver hepatocellular carcinoma (LIHC), and lung adenocarcinoma (LUAD), with positive hazard ratios indicating their potential role in promoting tumor progression (Figure 1B). There has been evidences indicating how METTL3/14 and IGF2BP1/3 have enhanced expression that stabilizes the expression of their targets and thus enhances cancer progression in an m6A-dependent manner, and therefore reduces overall survival in patients (N. Zhang et al. 2022; T. Li et al. 2019; Ding et al. 2025). Thus, these findings highlight the prognostic significance of m6A modification machinery in multiple cancers, where overexpression of both writers and readers may synergistically contribute to tumor progression and worse clinical outcomes.

Given the prominent upregulation of both m6A writers and readers in AML, we further analyzed the correlation between their expression levels in TCGA and TARGET AML datasets. METTL3 and METTL14 showed strong positive correlations with IGF2BP1, IGF2BP2, and IGF2BP3, and a moderate correlation with other components of the m6A machinery (Figure 1C).

In our previous studies, we established that IGF2BP1 and IGF2BP3 are overexpressed in ETV6-RUNX1 and MLL rearranged B-ALL, respectively, and increase the stability of their target genes (Sharma et al. 2021; Palanichamy et al. 2016) but the combined effect of m6A readers and writers on shared target genes has not yet been explored. To address this, we performed a proof-of-concept analysis integrating publicly available m6A-seq data from METTL3/METTL14 knockdown experiments and eCLIP-seq data for IGF2BP1/3, both generated in HepG2 cells. This helped us to identify the target genes that have m6A marks and are METTL3/14 dependent and are bound by IGF2BP1/3. The target genes were functionally validated to evaluate the combined regulatory effect of the m6A writers and readers on target gene expression, thereby shedding light on cooperative m6A-mediated post-transcriptional gene regulation in cancer.

### Intersection of m6A-Dependent IGF2BP1/3 Binding Sites Reveals Functionally Relevant Target Genes

To delineate the cooperative role of m6A writers (METTL3/14) and readers (IGF2BP1/3) in post-transcriptional gene regulation, we first identified target genes that are both methylated by METTL3/14 and bound by IGF2BP1/3. To identify the binding targets of the readers (IGF2BP1/3), the public eCLIP data for IGF2BP1 and IGF2BP3 for HepG2 cells were retrieved. The eCLIP-seq datasets of IGF2BP1/3 will contain the genomic intervals of the targets whose RNA was bound by IGF2BP1/3. The target genomic regions for IGF2BP1 and IGF2BP3 were identified and annotated using the optimized annotation pipeline (Figure 2a).

**Figure 2:**
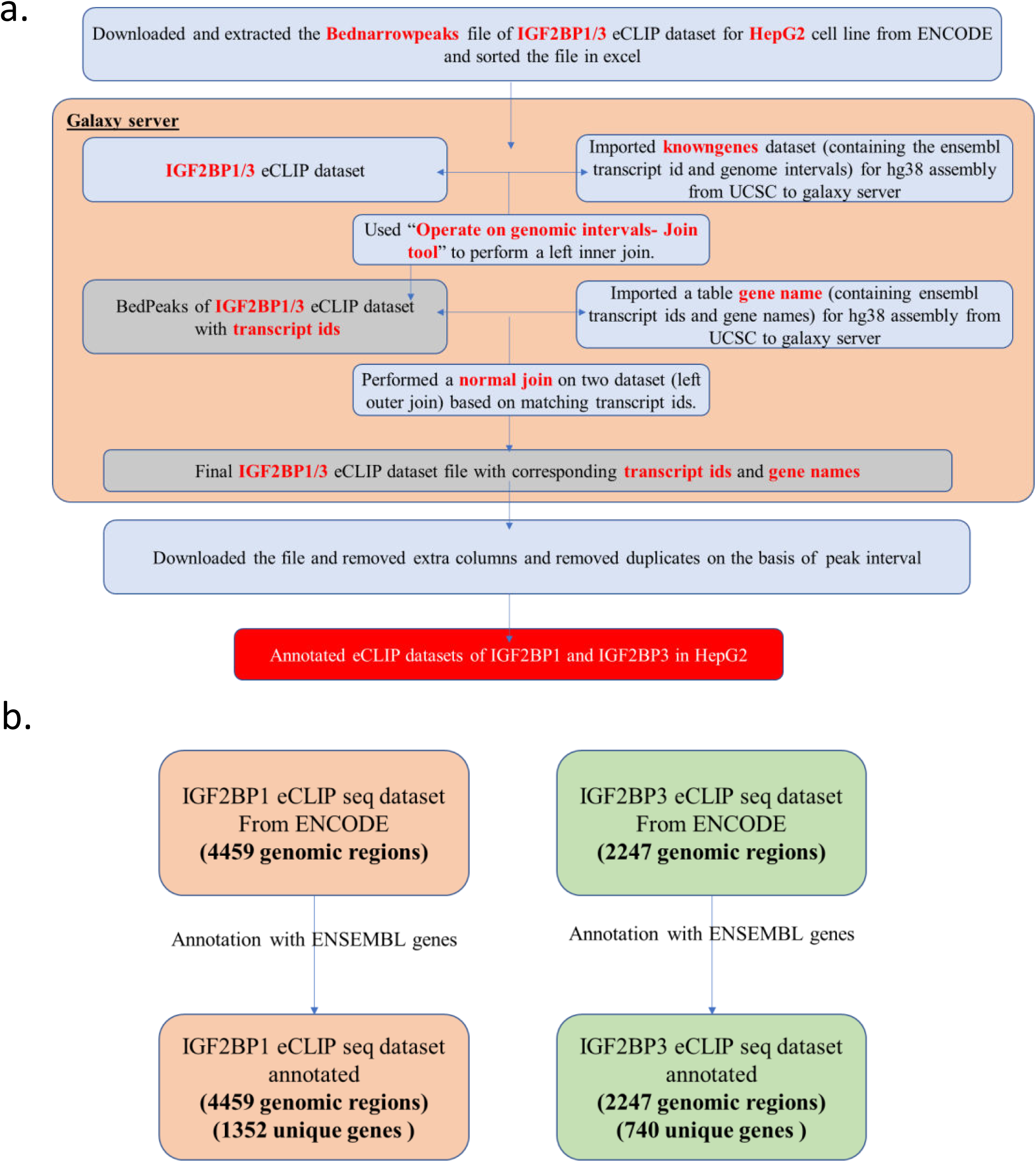
Annotation of pulled-down mRNAs with significant peak calling in IGF2BP1 and IGF2BP3 eCLIP-seq datasets from the public ENCODE repository. **a)** Pipeline for analysis and annotation of public eCLIP-seq datasets of IGF2BP1/3. **b)** Unique genes annotated from the public eCLIP-seq datasets of IGF2BP1/3.

In total, 4,459 genomic intervals corresponding to 1,352 unique genes were obtained for IGF2BP1, and 2,247 intervals representing 740 unique genes were identified for IGF2BP3. The difference between the number of genomic intervals and unique genes reflects the presence of multiple binding sites for each reader on a single transcript (Figure 2B).

To determine which of these reader-bound transcripts also require METTL3 or METTL14 for m6A deposition, we retrieved m6A-seq datasets from METTL3/14 knockdown (KD) experiments. These datasets include m6A-enriched transcriptomic regions in both control and METTL3/14 KD conditions. To identify the targets of readers that were dependent on writers for m6A methylation marks, we performed an intersection analysis of genomic interval files of both readers’ (annotated IGF2BP1/3 target genomic regions) and writers’ datasets (METTL3/14 m6A methylation-dependent targets). We defined METTL3/14-dependent methylation sites as those present in the control but lost upon METTL3 or METTL14 knockdown. These were identified by subtracting the methylated intervals in the KD condition from those in the control using our optimized analysis pipeline (Figures 3A, 3B).

**Figure 3:**
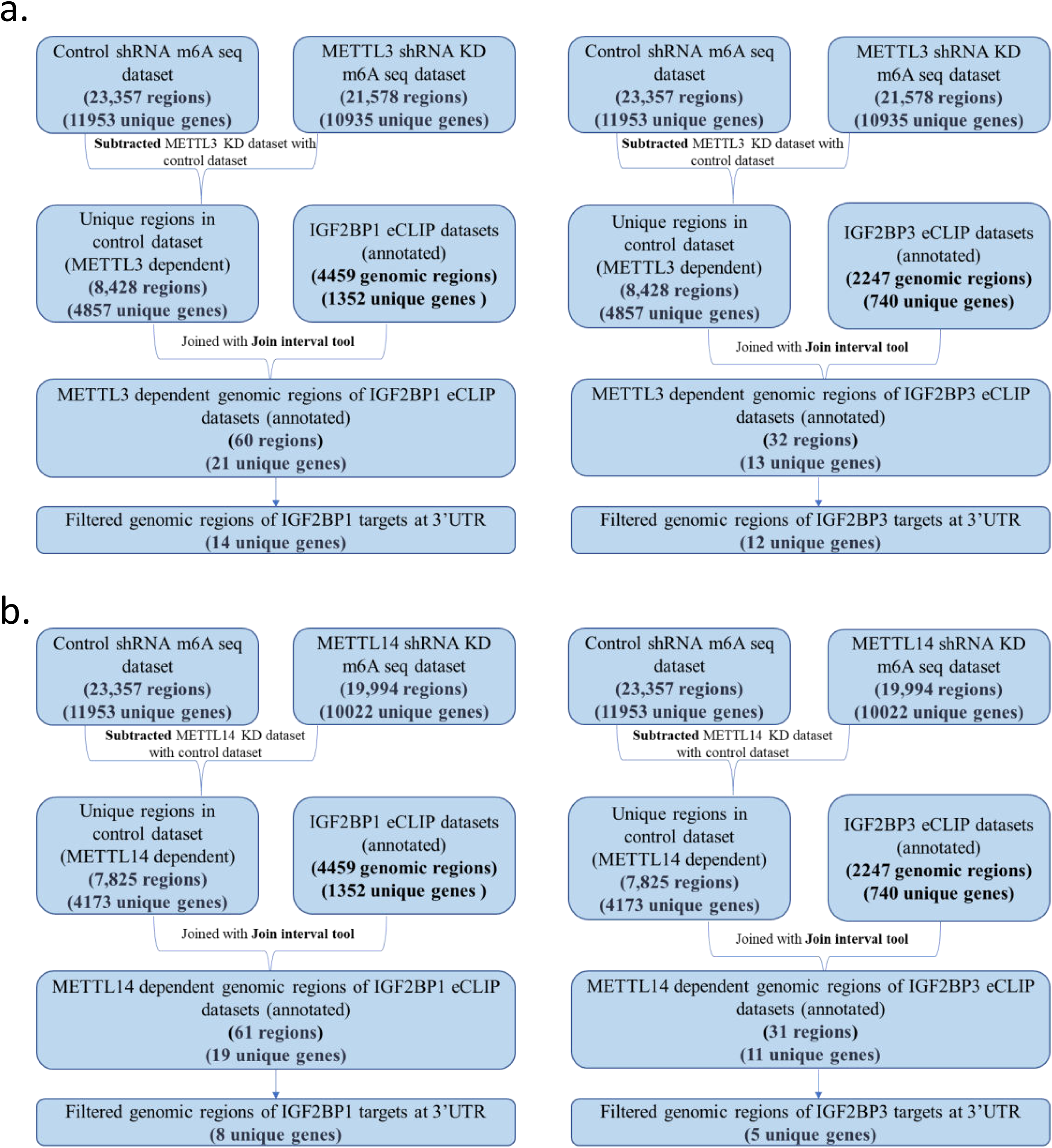
Deduction of m6A-dependent IGF2BP1 and IGF2BP3 eCLIP target genes from METTL3 and METTL14 knockdown datasets (GSE110320). The Analysis pipeline followed to identify intersection of IGF2BP1/3 eCLIP target genes with m6A-seq datasets after **a)** METTL3 knockdown and **b)** METTL14 knockdown

Next, we performed an intersection analysis between the annotated IGF2BP1/3 eCLIP peaks and the METTL3/14-dependent m6A regions. To further refine our search for functionally relevant targets, we focused on overlapping peaks located within the 3’-untranslated regions (3’-UTRs), as both METTL3/14 and IGF2BP1/3 are known to preferentially act through 3’-UTR methylation in regulating mRNA stability. 14 unique METTL3 dependent genes bound by IGF2BP1 and 12 unique genes were bound by IGF2BP3 were identified (Figure 3a, Table 1). These genes have m6A marks in their 3’UTR dependent on METTL3 and are also bound by IGF2BP1/3. 11 genes were found to be common targets of IGF2BP1 and IGF2BP3. Moreover, 8 unique genes were found for IGF2BP1, and 5 unique genes were found for IGF2BP3, which were METTL14 dependent (Figure 3b, Table 1). These genes have m6A marks in their 3’UTR dependent on METTL14 and are also bound by IGF2BP1/3.

**Table 1:** List of m6A-dependent IGF2BP1/3 eCLIP target genes identified from the intersection of the m6A-seq and IGF2BP1/3 eCLIP public datasets along with the oncogenic role, cancers associated, and biological functions. Target genes highlighted with green were taken forward for experimental validation.

| Gene | Role | Cancers Associated | Biological Function | m6A Dependence | eCLIP Target |
| --- | --- | --- | --- | --- | --- |
| DYNLL1 | Oncogene | Breast cancer | Regulates apoptotic activities of BCL2L11 by sequestering it to microtubules. | METTL14 | IGF2BP1/<br>IGF2BP3 |
| KDM3B | TSG | Colorectal cancer, leukemia (AML), prostate cancer, hepatocellular carcinoma | Demethylation of histone. | METTL14 | IGF2BP1 |
| RPL29 | Oncogene | Colon cancer, squamous cell carcinoma, pancreatic cancer | Cadherin binding, heparin binding. | METTL14 | IGF2BP1/<br>IGF2BP3 |
| SLC39A14 | Not Known | Colorectal cancer, prostate cancer, renal cell, gastric cancer | Modifies the activity of zinc-dependent phosphodiesterases. | METTL3/<br>METTL14 | IGF2BP1/<br>IGF2BP3 |
| SYNJ2 | Oncogene | Colorectal, prostate cancer | Mediates the inhibitory effect of Rac1 on endocytosis. | METTL3/<br>METTL14 | IGF2BP1 |
| GNA11 | Not Known | Uveal melanoma, breast cancer, and granulomas | Acts as an activator of phospholipase C. | METTL3 | IGF2BP1 |
| MARCHF8 | Not Known | Not known | E3 ubiquitin-protein ligase that mediates ubiquitination of CD86 and MHC class II. | METTL3 | IGF2BP1 |
| POLR2E | Not Known | Breast cancer, cervical, prostate cancer, thyroid cancer, AML | The structural component of Pol II, POLR2E/RPB5. | METTL3 | IGF2BP1/<br>IGF2BP3 |
| PSMG4 | Not Known | Indirect target of p50 and Bcl3 | Chaperone protein, which promotes assembly of the 20S proteasome. | METTL3 | IGF2BP1/<br>IGF2BP3 |
| REXO1 | Not Known | Not known | Exonuclease activity. | METTL3 | IGF2BP1/<br>IGF2BP3 |
| SLC22A23 | Not Known | Squamous cell carcinoma, TNBC | Transmembrane transporter activity | METTL3 | IGF2BP1/<br>IGF2BP3 |
| SOX12 | Not Known | Hepatocellular carcinoma, adenocarcinoma, colorectal, ovarian | Cell survival. | METTL3 | IGF2BP1/<br>IGF2BP3 |
| SREBF2 | Not Known | Oesophageal squamous cell carcinoma, TSG in prostate cancer | Regulates transcription of genes related to cholesterol synthesis pathway. | METTL3 | IGF2BP1/<br>IGF2BP3 |
| TRIM71 | TSG | Ovarian cancer | E3 ubiquitin-protein ligase. | METTL3 | IGF2BP1/<br>IGF2BP3 |
| MXD4 | Not Known | Ovarian cancer | Transcriptional repressor. | METTL3/<br>METTL14 | IGF2BP1/<br>IGF2BP3 |
| HNF4A | Not Known | Hepatocellular carcinoma, colon, rectal, and pancreatic cancer | Transcriptional regulator. | METTL3 | IGF2BP3 |
| CNOT1 | Oncogene | Osteosarcoma, hepatocellular carcinoma | Acts as a transcriptional repressor. | METTL14 | IGF2BP1 |
| PWP1 | Not Known | Pancreatic adenocarcinoma, leukemia | Regulates Pol I-mediated rRNA biogenesis and Pol III-mediated transcription. | METTL14 | IGF2BP1 |
| KANSL1 | Not Known | Not known | Acetylation of nucleosomal histone H4 and involved in transcriptional regulation. | METTL14 | IGF2BP3 |

The METTL3-dependent IGF2BP1/3 targets; *GNA11, SREBF2,* and *HNF4A* genes and METTL14-dependent IGF2BP1/3 targets; *CNOT1, DYNLL1, RPL29*, and *KDM3B* were selected to study the combined effect of IGF2BP1/3 and METTL3/14 on their target genes.

### METTLs and IGF2BPs have a combinatorial effect on their target genes via m6A marks on the 3’UTR

To investigate the combinatorial regulatory effects of m6A writers (METTL3/14) and readers (IGF2BP1/3) on specific target genes, we employed a luciferase reporter assay system. The 3′ untranslated regions (3′UTRs) of selected target genes-*GNA11, HNF4A, and SREBF2* (METTL3-dependent) and *CNOT1, KDM3B, RPL29*, and *DYNLL1* (METTL14-dependent)—were cloned downstream of the firefly luciferase gene in the pmirGLO dual-luciferase reporter vector (Figure 4A, S2, S3, S4). The cloned regions included sequences enriched in GGAC motifs and overlapping with IGF2BP1/3 binding peaks identified in eCLIP datasets (Figure S2), ensuring inclusion of functionally relevant m6A-marked regions.

**Figure 4:**
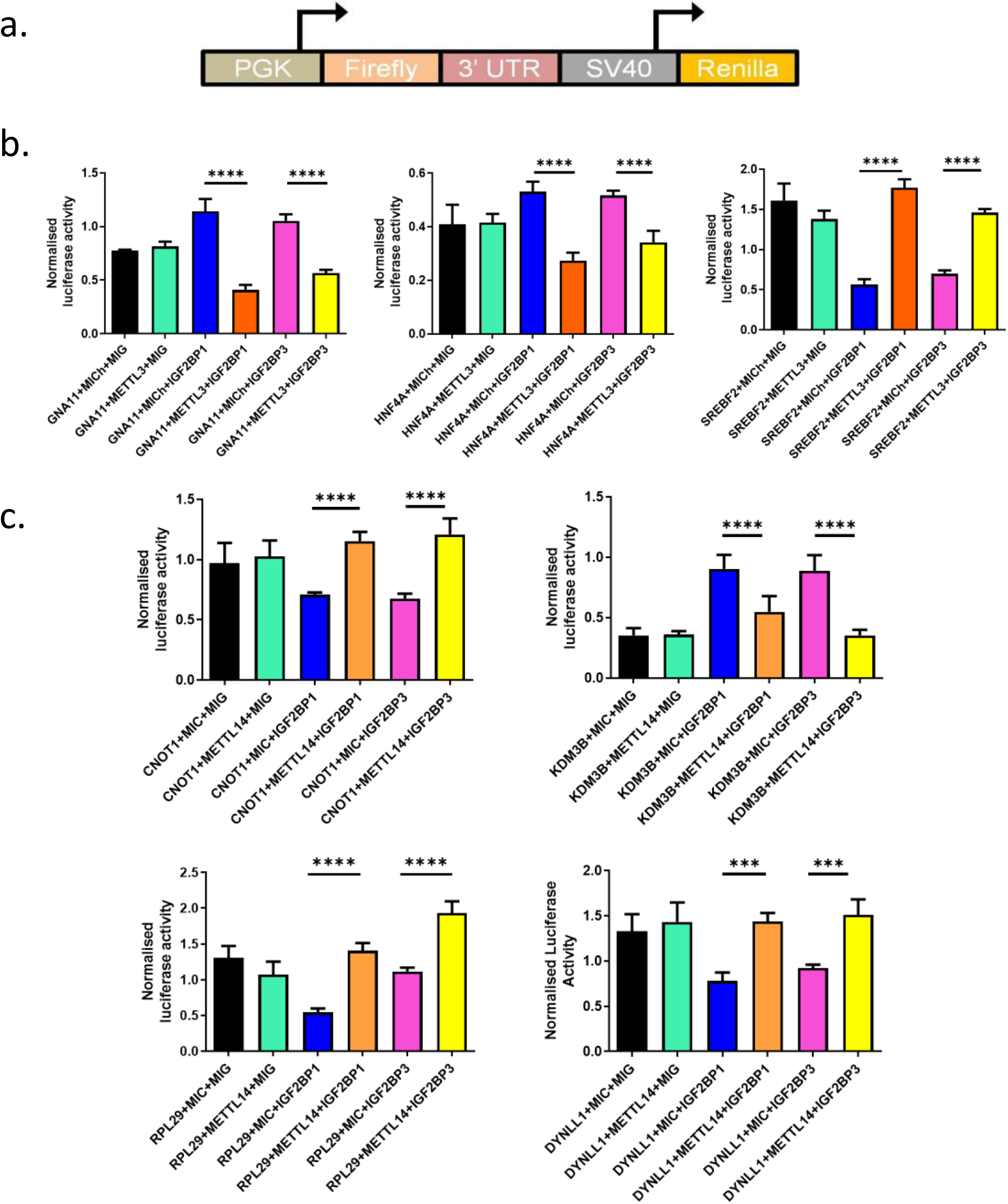
Effect of m6A writers (METTL3 and METTL14) and readers (IGF2BP1 and IGF2BP3) on their target gene expression via 3’UTR. a) Schematic of the 3’UTR-pmirGLO constructs of target genes generated for validation using the dual luciferase reporter assay. Dual-Luciferase reporter assay showing the combined effect of METTLs and IGF2BPs on **b)** METTL3 dependent and **c)** METTL14 dependent IGF2BP1/3 eCLIP targets. The dual luciferase activity was normalized to its respective empty control vector. The experiments were repeated thrice, and similar trends were observed. Graphs plotted show mean±SD (*t-test p<0.0001*).

To study the individual and combinatorial effect of METTLs and IGF2BPs on their targets, co-transfections were performed in HEK293T cells for transient overexpression of IGF2BP1, IGF2BP3, METTL3, and METTL14 along with the respective 3’UTR-Firefly luciferase constructs (Figure S5). The normalized Firefly/Renilla luminescence ratios (normalized luciferase activity) were analyzed for various combinations.

### METTL3 Targets: Differential Effects on IGF2BP-Mediated Stabilization

For *GNA11* and *HNF4A*, overexpression of IGF2BP1/3 alone led to a significant increase in luciferase activity, consistent with their role as mRNA stabilizers. Interestingly, METTL3 overexpression alone had no significant effect. However, co-expression of METTL3 with IGF2BP1/3 resulted in a significant reduction in luciferase activity compared to IGF2BP1/3 alone (Figure 4B). This suggests that METTL3-dependent methylation of the 3′UTR may interfere with IGF2BP1/3 binding or alter the stability of the transcript, ultimately reversing the stabilizing effect of the readers.

In contrast, *SREBF2* displayed a distinct regulatory pattern. While IGF2BP3 and METTL3 alone had no significant impact, IGF2BP1 alone reduced luciferase activity. Co-expression of METTL3 with IGF2BP1/3 led to a significant increase in luciferase activity, indicating that METTL3-generated m6A marks act as a stabilization signal for IGF2BP1/3 binding in this context. Thus, METTL3 and IGF2BP1/3 appear to synergistically enhance *SREBF2* expression.

### METTL14 Targets: Cooperative Stabilization with IGF2BP1/3

The METTL14-dependent targets *CNOT1, RPL29,* and *DYNLL1* exhibited increased luciferase activity upon overexpression of either METTL14 or IGF2BP1/3 alone. This effect was further amplified when both METTL14 and IGF2BP1/3 were co-expressed (Figure 4C), suggesting that m6A methylation by METTL14 facilitates IGF2BP1/3 binding and enhances mRNA stabilization. These results support a model in which METTL14 and IGF2BP1/3 cooperatively upregulate target gene expression.

Interestingly, *KDM3B* showed an inverse regulatory pattern. Luciferase activity was significantly reduced when both METTL14 and IGF2BP1/3 were co-expressed, whereas overexpression of either component alone led to an increase in reporter activity. These findings suggest that METTL14-dependent m6A marks may serve as destabilization signals in the context of *KDM3B*, and that IGF2BP1/3 may mediate transcript degradation or repress translation when guided by such marks.

## Discussion

N6-methyladenosine (m6A) is the most prevalent internal, co-transcriptional modification in eukaryotic mRNA, predominantly occurring within the consensus RRACH motif and frequently enriched in the 3′ untranslated region (3′UTR) of transcripts. The m6A modification machinery consists of three main classes of proteins: writers, erasers, and readers. Writers, such as METTL3 and METTL14, catalyze the methylation reaction; erasers like ALKBH5 and FTO remove m6A marks; and readers, including IGF2BP1/3, are RNA-binding proteins that recognize methylated transcripts and influence downstream outcomes (Fang et al. 2022; Wang et al. 2022). The methylated mRNA is recognized by various mRNA-binding proteins, which can result in mRNA stability by increasing its half-life and further promoting translation (Balacco and Soller 2019; Wu et al. 2016; Weng, Huang, and Chen 2019). The WERs of m6A machinery have been known to act as oncogenes in various cancers.

Aberrations in m6A regulators have been widely implicated in oncogenesis across multiple cancers, including breast, lung, colorectal, liver, leukemia, glioblastoma, kidney, and pancreatic cancer (Fang et al. 2022; Gao et al. 2021). Depending on the cellular and tumor context, components of the m6A machinery may function as oncogenes or tumor suppressors. Previous studies from our group demonstrated that IGF2BP1 is overexpressed in ETV6::RUNX1-positive B-ALL, and IGF2BP3 in MLL-rearranged B-ALL, where they act to stabilize oncogenic transcripts (Sharma et al. 2021; Palanichamy et al. 2016). Pan-cancer analysis using the public NCBI-GEO, TCGA, and TARGET datasets showed dysregulated expression of WERs in different cancer datasets (Figure 1a, SF1). Moreover, the correlation analysis in AML, LIHC, and LUAD datasets showed an inverse correlation between writers’ and readers’ expression with patient survival (Figure 2a). However, the cooperative interplay between m6A writers and readers, especially in terms of their effect on shared targets, remains incompletely understood.

In this study, we systematically explored the combined effect of IGF2BP1/3 (readers) and METTL3/14 (writers) on downstream gene regulation. Using integrative analyses of m6A-seq datasets (following METTL3/14 knockdown) and eCLIP-Seq datasets for IGF2BP1/3, we identified a set of high-confidence shared target genes that are methylated by METTL3 or METTL14 at their 3′UTRs and are directly bound by IGF2BP1/3. These targets were subsequently validated using dual-luciferase reporter assays, where the 3′UTRs of selected genes were cloned downstream of a luciferase reporter to assess the post-transcriptional effects of reader and writer overexpression.

Our results revealed two distinct regulatory patterns:

### Synergistic Stabilization

For several targets, including *SREBF2* (METTL3-dependent), *RPL29, DYNLL1*, and *CNOT1* (METTL14-dependent), co-expression of METTL3 or METTL14 with IGF2BP1/3 led to significantly increased luciferase activity compared to either factor alone. These findings support a model in which methylation by METTL3/14 enhances IGF2BP1/3 binding and leads to transcript stabilization. The writers produce m6A marks on the 3’ UTR of the gene, which enhances the binding of IGF2BPs and ultimately increases the expression of the gene, as shown by *c-MYC* (Sharma et al. 2021). This is consistent with previous studies where IGF2BP3 stabilized oncogenic transcripts such as *PD-L1* in breast cancer (Wan et al. 2022) and spermine synthase (SMS) in pancreatic cancer (Guo et al. 2022), both through m6A-dependent binding. This further leads to the disruption of immune surveillance (Wan et al. 2022) and regulation of PI3K-AKT/EMT signaling pathways and thus enhancing pancreatic cancer progression (Guo et al. 2022). Similarly, IGF2BP1/3 may recognize methylated regions and promote mRNA stability, enhancing translational output and potentially promoting oncogenesis. This supports the notion that m6A methylation and reader recognition can function cooperatively to upregulate gene expression.

### Antagonistic Destabilization

In contrast, other targets such as *HNF4A* and *GNA11* (METTL3-dependent), and KDM3B (METTL14-dependent) demonstrated an opposite trend. While IGF2BP1/3 alone increased luciferase activity, consistent with their stabilizing role, co-expression with METTL3 or METTL14 resulted in reduced activity, suggesting a destabilizing effect. This implies that in some cases, m6A methylation may hinder reader binding or recruit destabilizing cofactors. A possible explanation is that methylation within certain regions of the 3′UTR alters RNA secondary structure or accessibility, reducing IGF2BP affinity. The m6A-containing mRNA is destabilized by YTHDF2, an RNA-binding protein that recruits the CCR4-NOT deadenylase complex, which deadenylates m6A-containing mRNA and thus leads to its destabilization and decay. (Du et al. 2016) IGF2BP1 interacts with long noncoding RNAs like *HULC* and UCA1 and destabilizes them. Moreover, IGF2BP1 recruits the CCR4-NOT1 deadenylase complex, which leads to its destabilization and degradation (Zhou et al. 2018; Hämmerle et al. 2013; Duan et al. 2024). Alternatively, the position and context of m6A marks, or the sequence architecture of the 3′UTR, may dictate whether IGF2BP1/3 function as stabilizers or destabilizers. These findings highlight a context-dependent role of m6A readers and raise the possibility of dual functionality, acting either to stabilize or destabilize transcripts depending on sequence context and the specific interplay with writer activity (Figure 5).

**Figure 5:**
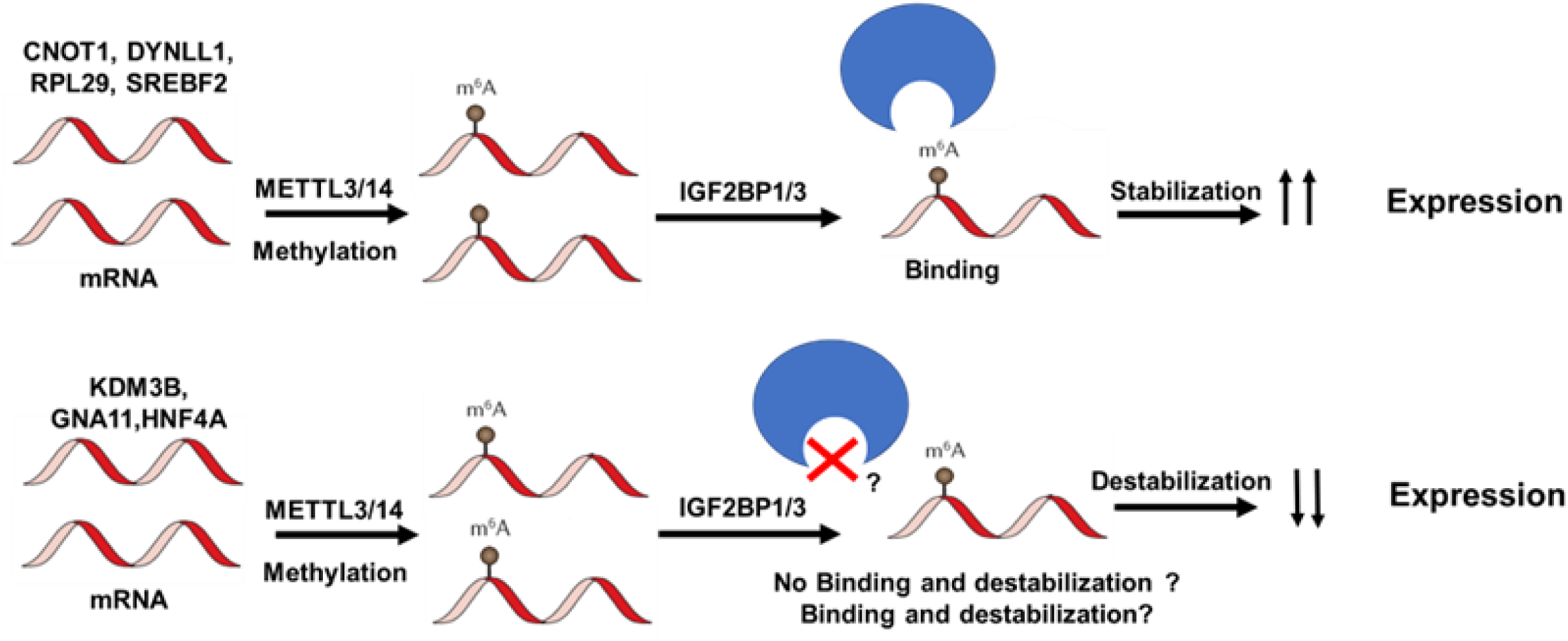
Graphical summary of the study. The proof-of-concept study of the combined effect of m6A writers (METTL3 and METTL14) and readers (IGF2BP1/3) on the expression of their target genes.

Notably, previous transcriptomic studies using IGF2BP1/3 knockdown followed by RNA-seq have shown that while many target genes are downregulated (suggesting a stabilizing role), others are upregulated, implying that IGF2BP1/3 may also promote mRNA degradation or repress translation in certain contexts (Palanichamy et al. 2016; Elcheva et al. 2020). Our findings for *HNF4A, GNA11*, and *KDM3B* align with this possibility and provide mechanistic insight into this underexplored aspect of IGF2BP biology.

Taken together, our study provides the first experimental evidence of how m6A writers and readers can exert combinatorial effects on shared target genes. Importantly, these effects are not uniform; instead, they are target-specific, with some genes exhibiting enhanced stability and others showing destabilization in response to methylation and reader interaction. This suggests a nuanced and varied regulatory landscape where m6A modifications serve as tunable signals that modulate RNA fate in a gene-specific manner.

## Conclusion

This proof-of-concept study establishes that the m6A writers METTL3 and METTL14 and readers IGF2BP1 and IGF2BP3 have a combined, yet context-dependent effect on their target mRNA stability, which could be either stabilization or destabilization, and this effect is target-specific. These effects may be synergistic, leading to increased gene expression, or antagonistic, resulting in transcript destabilization and reduced expression. Our work supports a broader paradigm in which m6A marks function as regulatory switches, integrating writer and reader activity to fine-tune gene expression outcomes.

Further studies are needed to understand the structural basis of reader binding specificity, the role of other cofactors, and the *in vivo* relevance of these interactions in different cancer models. Understanding how this dynamic is influenced by therapeutic interventions may offer new avenues for targeting the m6A machinery in cancers characterized by dysregulated post-transcriptional gene regulation.

## Material and Methods

### Data acquisition and analysis from TCGA database

Pan-cancer analysis of m6A machinery proteins was conducted using publicly available datasets from NCBI-GEO, TCGA, and TARGET, along with control datasets from the GTEx repository. For pan-cancer multigene expression dot matrix comparisons of writers, erasers, and readers (WERs), data from NCBI-GEO, TCGA, TARGET, and tissue-matched GTEx datasets were retrieved from the TNMplot webserver (Bartha and Győrffy 2021). The TNMplot web server facilitates multiple gene expression data comparisons between multiple groups of tumor studies and normal samples to determine the log2 fold-change (FC) values. Survival analysis of m6A machinery genes in relation to overall survival, as well as gene-gene correlation matrices, was performed using the GEPIA2 web portal (http://gepia2.cancer-pku.cn/) (Tang et al. 2019). To perform these analyses, pan-cancer datasets from TCGA and TARGET studies including the ACC, BRCA, CESC, CHOL, COAD, DLBC, ESCA, GBM, HNSC, KICH, KIRC, LAML, LIHC, LUAD, LUSC, PRAD, READ, SARC, SKCM, STAD, TGCT, THCA, THYM, UCEC and UCS, along with tissue-matched GTEx datasets, were retrieved via the GEPIA2 portal.

### Genomic interval intersection analysis

This technique was performed to identify the common targets of the m6A writers and readers (IGF2BP1/3). The unique binding target mRNAs of the readers IGF2BP1 and 3 were identified using publicly available eCLIP datasets. The bednarrowpeaks files of eCLIP-Seq of IGF2BP1 (GSE92021) and IGF2BP3 (GSE92220) (The ENCODE Project Consortium 2012) for HepG2 cells were downloaded from ENCODE (https://www.encodeproject.org/) and then uploaded along with the hg38 assembly from UCSC on the Galaxy Server (https://usegalaxy.org). The two datasets, eCLIP-Seq data for IGF2BP1/3 and METTL3/14 KO datasets, were joined using the “Operate on genomic intervals-Join tool,” which helps join two datasets based on the overlapping genomic intervals. The left inner join was performed to record the genomic regions present in the IGF2BP1/3 eCLIP dataset. The bedpeaks of IGF2BP1/3 with transcript IDs were generated. The known gene dataset containing transcript IDs and gene names was imported from UCSC and was added using the “normal join” function. The data was annotated based on matching Ensembl transcript IDs.

The GSE110320 dataset has m6A-seq datasets from HepG2 cell line under three conditions: 1) Control 2) METTL3 knockdown 3) METTL14 knockdown (H. Huang et al. 2019). The database refers to genomic regions for target mRNA pulled down using the anti-m6A antibody. To identify the target genes dependent on METTLs, the genomic regions extracted from the shControl m6A seq dataset were subtracted from the METTL3 or METTL14 KD dataset using the “subtract whole dataset” function. This subtraction specifically identifies those genomic regions not present in the METTL3/14 KD dataset and thus depend on METTL3/14 for m6A methylation. These METTL3/14 dependent regions that were unique in the control dataset were further used for the downstream analysis.

The intersection of the annotated genomic intervals of these two datasets was performed. The annotated IGF2BP1/3 eCLIP dataset and METTL3/14 dependent regions were joined using the left inner join from the “operate on the genomic interval -join tool.” The gene structure file containing annotated genomic regions like exon, intron, 3’UTR, 5’UTR, gene names, and transcript IDs was imported from UCSC. The left inner join using the “operate on genomic interval-join tool” was performed on gene structure files and METTL3/14 and IGF2BP1/3 common region files. This gave us the list of METTL3/14 dependent regions bound by IGF2BP1/3 on 3’UTR. These targets were then further validated using luciferase assay.

### Plasmids and cloning of m6A machinery and target genes 3’UTR

The m6A writers METTL3/14 were cloned into the MSCV-IRES-mCherry (MICh) vector, and m6A readers IGF2BP1/3 were into the MSCV-IRES-GFP (MIG) vector between BglII and XhoI sites (Figure S3) (Table S1) downstream of the MSCV promoter. MSCV-METTL3/14-IRES-mCherry (MICh-METTL2/14) and MSCV-IGF2BP1/3-IRES-GFP (MIG-IGF2BP1/3) constructs and were used for the transient overexpression of readers and writers. The cloning was performed using the standard protocol as described previously (Palanichamy et al. 2016).

The 3’UTR of the selected genes were cloned in the pmirGLO vector (Figure S4) Cloning primers flanking the 3’UTR were designed for the target genes with overhangs of restriction sites to be inserted in the vector between SacI and XhoI as described previously (Palanichamy et al. 2016) (Table S1).

### Cell Culture and transfections

The HEK293T cell line was obtained from American Type Culture Collection (ATCC-CRL3216) and maintained in DMEM (Dulbecco’s Modified Eagle’s Medium, Gibco), supplemented with 10% (v/v) FBS (fetal bovine serum, Gibco), 2 mM L-Glutamine (Gibco) and 10μg/ml Pen-Strep (ThermoScientific) in a 5% CO_2_ incubator at 37 °C. The cells were cultured in 10 cm^2^ sterile culture plates and passaged every 3-4 days before reaching full confluency. The HEK 293T cells were plated in 24-well plates at 50,000 cells/well density and transfected at 70% confluency using PEI-Max (24765, Polysciences) transfection reagent.

### Luciferase assays

The plasmids carrying IGF2BP1/3 (MIG) and METTL3/14 (MICh), along with the empty pmirGLO/3’UTR constructs of target genes, were co-transfected in a 5:5:1 ratio (250:250:50 ng), respectively, into the HEK293T cells plated in a 24-well plate (Figure S5). After 48 hours, the cells were lysed using 100μL 1X Passive Lysis Buffer (PLB), and the cell lysates were centrifuged at 13,000g for 5 minutes to clear the cell debris.

The luciferase assay was done using the Dual-Luciferase Reporter Assay kit (Promega) as per the manufacturer’s instructions. Luminescence was measured using a GloMax luminometer (Promega). The ratios of firefly to Renilla luciferase activity were determined for all combinations. The ratio obtained from combinatorial co-transfections was normalized with empty vector control (MIG+MICh+pmirGLO). Each 3’UTR construct co-transfection combination had 3’UTR-pmirGLO plasmid along with either both IGF2BP1/3-MIG and METTL3/14-MICh or one empty vector control, i.e., either IGF2BP1/3-MIG or METTL3/14-MICh or both empty vector controls, i.e., MIG and MICh, and the normalized luciferase activity was compared across the combinations.

### Statistical analysis

All the experiments were performed in triplicate, and *in vitro* experiments were repeated thrice. Data represent mean ±SD for continuous numerical data. 2-tailed Student’s *t-*tests were performed using GraphPad Prism software and applied to each experiment as described in the figure legends. A *P* value less than 0.05 was considered significant. \**P* < 0.05, \*\**P* < 0.01, \*\*\**P* < 0.001, and \*\*\*\**P* < 0.0001.

## Author Contributions

RB and HB conducted the Bioinformatics analysis and molecular biology experiments and analyzed the data. RB, HB, and JKP wrote the manuscript. JKP conceived the study. All authors reviewed and edited the manuscript.

Conceptualization; R.B., H.B., and J.K.P.; Methodology, R.B., H.B.; Validation, R.B., H.B., and J.K.P.; Formal analysis; Investigation, R.B., H.B., and J.K.P.; Data curation, R.B., H.B; Writing— original draft preparation, R.B., H.B.; Writing—review and editing, R.B., H.B., and J.K.P.; Visualization; R.B., H.B., Supervision J.K.P.; Project administration, R.B., H.B., and J.K.P.; Funding acquisition, J.K.P. All authors have read and agreed to the published version of the manuscript.

## Funding and Acknowledgments

This work was supported by a SERB Core Research Grant (CRG/2021/004251) and an AIIMS-THSTI Collaborative Intramural Grant to JKP. RB is supported by a DBT Junior Research Fellowship, and HB is supported by a DBT Senior Research Fellowship.

R.B. is thankful to the Department of Science and Technology-Science and Engineering Research Board (DST-SERB) for the Junior Research Fellowship (March 2022 -April 2023) and Department of Biotechnology, Government of India, for the Junior Research Fellowship (May 2023-Present). H.B. is thankful to the Department of Biotechnology, Government of India for Junior Research Fellowship (December 2021–December 2023) and Senior Research Fellowship (December 2023–Present).

## Institutional Review Board Statement

The study was reviewed and approved by the Institutional Ethics Committee of All India Institute of Medical Sciences (Ref. No.: IECPG-51/27.02.2020, RT-29/26.08.2020, IECPG57/27.02.2020, RT-29/26.08.2020 and date of approval 29.08.2020).

## Data Availability Statement

The paper and supplementary information contain all the data to support the study’s findings. Any additional information required will be provided on request.

